# Programming Primary Schwann Cells with Circular RNA Delivered by *Listeria* Drones

**DOI:** 10.64898/2026.11.10.765776

**Authors:** Keit Men Wong, Oliver Trunschke, Kiet N. Tran, Mark Gomelsky, Jared S. Bushman

## Abstract

Efficient transient genetic engineering of primary cells remains a major challenge in regenerative medicine, immunotherapy, and other cell-based therapies, as many therapeutically relevant cell types are refractory to conventional gene delivery. Primary Schwann cells (SCs) hold particular promise for peripheral nerve and spinal cord regeneration, yet nonviral gene delivery into these cells remains challenging. Here, we demonstrate transient programming of primary rat SCs using circular RNA (circRNA) delivered by attenuated, self-destructing *Listeria monocytogenes* (*Lm*) drones. These engineered bacteria invade mammalian cells, escape the phagosome, synthesize and circularize target RNA under inducible control, and undergo programmed lysis to release circRNA into the host cytosol for IRES-mediated protein synthesis. Previously validated in cancer and immune cell lines, this platform had not been evaluated in primary cells. Using a reporter nano-luciferase (NLuc)-encoding circRNA, we optimized SC bactofection across multiplicities of infection (MOIs). At MOIs of 10-20, bactofection efficiency reached 94%, with negligible SC death. Viable *Lm* were completely eliminated through programmed bacterial lysis and antibiotic treatment, yielding bacteria-free SCs that continued expressing the NLuc reporter for over five days and were suitable for transplantation. The platform enables straightforward replacement of NLuc with one or more therapeutic payloads. These findings establish the first demonstration of *Lm* drone-mediated circRNA delivery into primary cells. By enabling transient, non-integrating expression of therapeutic factors in difficult-to-transfect SCs, this approach offers a promising strategy for developing temporally controlled cell therapies for peripheral nerve and spinal cord repair.

## 1. Introduction

Efficient engineering of primary cells remains a major challenge in regenerative medicine, immunotherapy, and other cell therapies because many therapeutically relevant cell types are refractory to conventional gene delivery methods. Primary Schwann cells (SCs) represent a particularly challenging example owing to their low transfection efficiency and sensitivity to physical and chemical stressors [1–6]. At the same time, SCs are attractive therapeutic candidates because they promote axonal regeneration, remyelination, and trophic support following peripheral nerve injury and have also been investigated as cellular therapeutics for central nervous system repair [7–11]. Because the regenerative phenotype of SCs is naturally transient, strategies that enable temporary rather than permanent genetic programming are particularly attractive. However, efficient methods for transient intracellular engineering of primary SCs remain lacking.

Conventional non-viral approaches, including lipofection and nucleofection, generally produce low transfection efficiencies while causing cell aggregation, cellular stress, and reduced viability (Table 1) [1–6]. Viral vectors, including lentiviral and adeno-associated viral (AAV) systems, improve gene transfer efficiency [12–15] but require specialized vector production, are constrained by cargo capacity, and typically result in persistent transgene expression [16]. Although these approaches are valuable in some settings, transient protein expression is preferable for applications requiring temporary modulation of cell phenotype. Thus, there is a need for alternative intracellular delivery strategies that enable transient genetic programming without compromising SC viability.

**Table. 1.** Transfection efficiencies of nonviral delivery methods in primary SCs.

| Method / reagent | SC source | Cargo | Efficiency | Ref. |
|---|---|---|---|---|
| Self-destructing *Lm* drones | Rat | circRNA | 94% | This work |
| Lipofection (FuGENE HD) | Rat | Plasmid | 2% | [1] |
| Nucleofection (AMAXA) | Rat | Plasmid | 10% | [1] |
| Electroporation / nucleofection | Human/rat | Plasmid | 38–40%<br>20-33% | [2] |
| Polymer (Viromer RED) | Rat | Plasmid | 20% | [3] |
| Lipofection (Lipofectamine 3000) | Rat | Plasmid | 20% | [3] |
| Nucleofection (Nucleofector I) | Human | Plasmid | 40–60% | [4] |
| Lipofection (reagents unspecified) | Human | Plasmid | < 5% | [4] |
| Lipofection 2000 or nucleofection II | Rat | Plasmid | 15–20% | [5] |
| Calcium phosphate co-precipitation | Rat | Plasmid | 0.01% | [6] |

*Listeria monocytogenes* (*Lm*) offers an alternative intracellular delivery strategy because it naturally invades mammalian cells, escapes from the phagosome, and accesses the host-cell cytosol [17, 18]. These properties have been exploited for bacterial-mediated nucleic acid delivery ("bactofection") using highly attenuated *Lm* strains, yet efficiency of gene delivery was modest [19–24]. We recently re-invigorated this concept by engineering self-destructing *Lm* "drones" that combine intracellular bacterial invasion with programmable synthesis and release of circular RNA (circRNA). In this platform, intracellular *Lm* synthesize RNA under the anhydrotetracycline (aTc)-inducible T7 Polymerase-driven promoter, efficiently circularize transcripts using a self-splicing ribozyme derived from the *Tetrahymena thermophila* group I intron, and subsequently undergo programmed lysis due to the phage-derived holin and lysin system. As result of the lysis, intracellular *Lm* releases circRNA directly into the host-cell cytosol where it can be translated by the host ribosomes via the internal ribosome entry site (IRES)-mediated translation (Fig. 1A). Unlike conventional RNA delivery approaches that require purification and intracellular delivery of exogenous RNA, the platform generates circRNA immediately before its release into the host cytosol. By integrating intracellular RNA synthesis, circularization, and cytosolic release within a single biological chassis, the platform eliminates multiple manufacturing and delivery steps required for conventional RNA therapeutics while allowing diverse protein programs to be implemented simply by exchanging the circRNA expression cassette. Because only the circRNA payload is exchanged, the *Lm* chassis offers a modular system that may be adaptable to diverse primary cell types (Fig. 1A).

**Figure 1.**
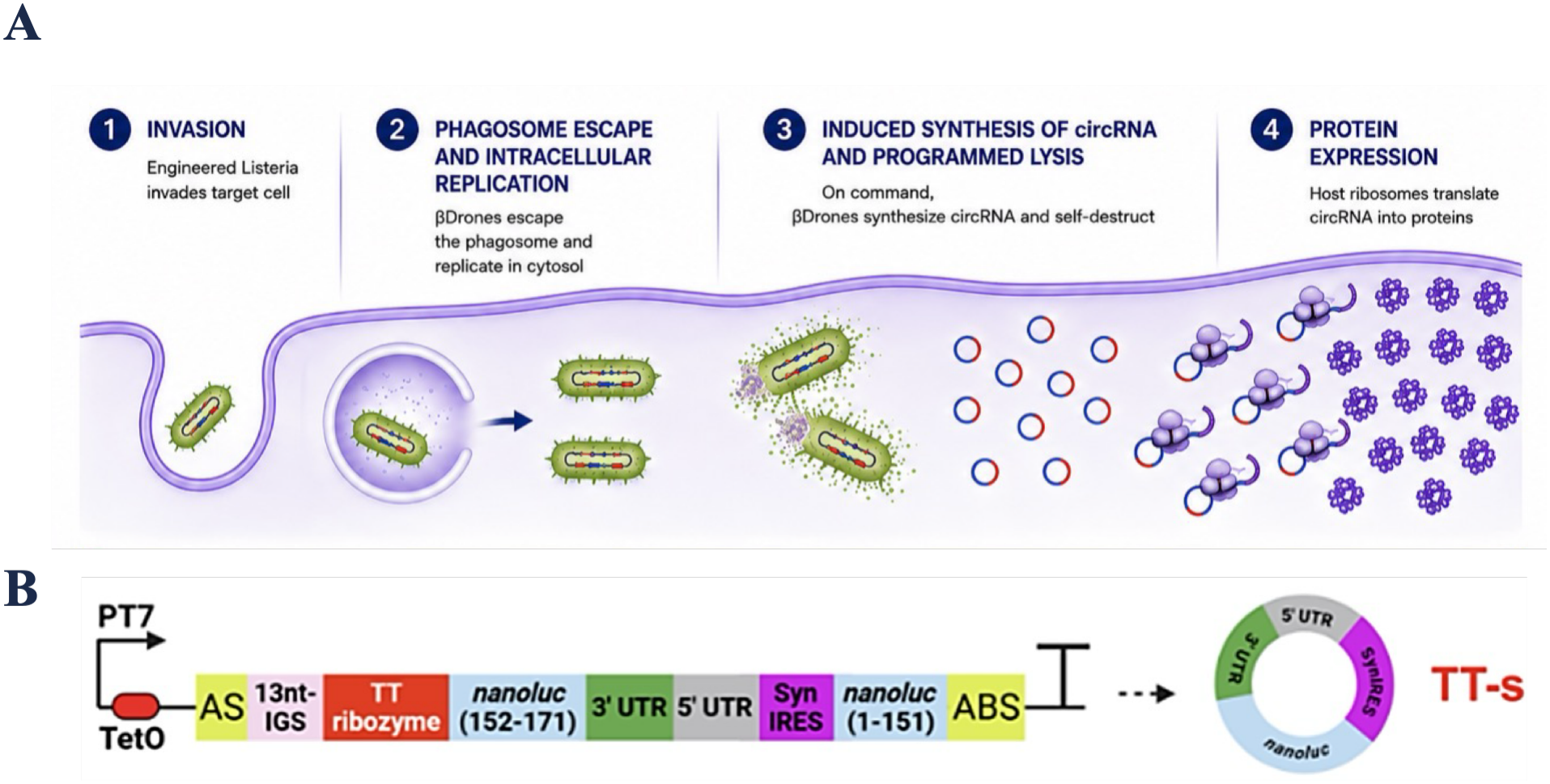
*Lm* drone-mediated Intracellular circRNA Delivery for Primary Schwann Cell Engineering. **A.** Schematic of the *Lm*-circRNA delivery platform. (1) *Lm* invades primary SCs, (2) escapes the phagosome, and (3) produces circRNA under aTc-inducible control. (3) aTc-induced holin–lysin-mediated bacterial lysis releases the circRNA cargo into the SC cytosol, where (4) IRES-driven translation produces the NLuc protein. **B.** aTc-inducible TT-s split-NLuc circRNA construct used in primary SCs. The construct contains the aTc-inducible T7 Polymerase driving expression of the TT-s circRNA cassette, including the internal guide sequence, TT ribozyme, split NLuc fragments (amino acids 152–171 and 1–151), 5′ and 3′ untranslated regions (UTRs), and synthetic IRES; the corresponding circularized RNA product is shown at right.

Circular RNA is a covalently closed structure, which greatly increases its resistance to exonuclease-mediated degradation and supports prolonged protein expression [25–26]. IRESs enable cap-independent translation, allowing engineered circRNAs to function as protein-expression templates without conventional 5′ caps or poly(A) tails [27]. Together, intracellular circRNA synthesis and programmed bacterial lysis eliminate the need for RNA purification and formulation while providing a transient, non-integrating route to protein expression that complements existing RNA delivery technologies (Fig. 1B).

Using this strategy, we recently established an inducible *Lm*-drone circRNA platform that achieves up to 95% RNA circularization efficiency for small inserts and 80-200-fold greater protein expression than corresponding linear RNA constructs in multiple rodent and human cancer cell lines, including a rat Schwannoma cell line [28]. Whether this intracellular circRNA engineering strategy can be extended from immortalized cell lines to therapeutically relevant primary cells, however, remained unknown.

In this study, we evaluated whether the self-destructing *Lm* circRNA platform could be adapted for transient genetic engineering of primary rat SCs. Using our previously validated split-NLuc circRNA reporter, we characterized bacterial infection and intracellular localization, assessed inducible circRNA expression, and systematically optimized the multiplicity of infection (MOI) and post-infection timing to maximize delivery efficiency while minimizing cytotoxicity. We also characterized the kinetics and persistence of circRNA-driven reporter expression and optimized the protocol for complete bacterial clearance following antibiotic treatment. Because the platform enables straightforward replacement of the NLuc reporter with therapeutic payloads, these studies establish an optimized workflow for circRNA bactofection of primary rat SCs and define key parameters governing delivery efficiency, expression kinetics, cytotoxicity, and bacterial clearance. More broadly, our findings provide a foundation for extending this temporary intracellular engineering strategy to other difficult-to-transfect primary cell types.

## 2. Materials and Methods

### 2.1 Primary Schwann Cell Isolation and Purification

All procedures involving animals were approved by the Institutional Animal Care and Use Committee (IACUC) of University of Wyoming.

Primary SCs were isolated from postnatal day 3 (P3) Sprague-Dawley rat sciatic nerves and expanded as previously described [29, 30]. Briefly, dissected nerves were mechanically shredded and enzymatically dissociated (1% collagenase, Worthington LS004196; 0.25% trypsin, Gibco 15090-046; 30 min, 37°C), then plated on poly-L-lysine–coated dishes (1 × 10^6^ cells/100-mm dish) in DMEM with 10% fetal calf serum. Contaminating fibroblasts were suppressed with 10 μM Ara-C (Sigma C6645) for 48 h, and SCs proliferation was promoted with 2 μM forskolin (Sigma 344270) and 2 ng/mL RhNRG (R&D Systems 396-HB) beginning ∼day 6–7. Residual fibroblasts were eliminated by complement-mediated cytolysis using anti-Thy1.1 IgM (Bio-Rad MCA04G) and rabbit complement (Sigma), repeated as needed. Purified SCs were expanded through two passages and cryopreserved at passage 3. SC purity was assessed by immunofluorescence staining for the SC marker SOX10, with purity exceeding 95% across passages 1–3 (Supplementary Figure 1). All primary SCs experiments were performed using cells below passage 10.

### 2.2 Bacterial Strains and Construct Design

An attenuated strain *Lm* XFL-7 constitutively expressing GFP and carrying the previously described and validated aTc-inducible TT-s circRNA system was used throughout this study [28]. The construct contains a split-NLuc reporter that generates functional luminescence only following proper splicing and precise mRNA circularization. Proper circularization of the circRNA was previously confirmed by RNase R resistance and sequencing across the predicted splice junction [28]. The delayed aTc-inducible holin–lysin lysis system is encoded in the same strain [28]. *Lm* was cultured in Brain-Heart Infusion (BHI) broth at 37 °C with appropriate antibiotic selection, and bacterial density was determined by OD_600_ for preparation of infection inocula.

### 2.3 circRNA Bactofection Protocol

SCs were seeded in a 96-well plate the night before infection and grown to ∼80% confluency prior to infection. On the day of infection, *Lm* cultures were washed in DMEM, resuspended in antibiotic-and serum-free DMEM, and added to SCs at indicated multiplicity of infection (MOI) of 5–100 for 1 hour (t = 0 h, the moment *Lm* was added to the culture). Extracellular bacteria were removed by washing, followed by incubation in medium containing 2% FBS and 50 µg/mL gentamicin for 1 hour, which ensures elimination of unwashed extracellular *Lm*. Cells were subsequently maintained in SC complete growth medium containing gentamicin until aTc induction and/or assessment at indicated time points. Induction was performed with 1 µg/mL aTc for 6 to 24 h. Unless otherwise indicated, moxifloxacin (4 µg/mL) was added following 6 or 24 h of aTc induction and maintained for the indicated duration to clear intracellular *Lm* bacteria.

### 2.4 Assessment of Intracellular Localization and Infection Efficiency

To evaluate the efficiency of Lm infection of primary SCs, intracellular bacteria were assessed as follows. SCs were infected at an MOI of 20 as described, fixed with 4% paraformaldehyde 24 h post-infection, permeabilized, and co-stained with DAPI (nuclei) and an anti-SOX10 antibody (SC identity). Fluorescence images were acquired on an EVOS FL microscope (Thermo Fisher). Infection efficiency (%) was quantified in ImageJ as the number of GFP⁺ cells (GFP serving as an *Lm* marker) divided by the total number of DAPI⁺ nuclei and expressed as a percentage.

### 2.5 SC Viability Assessment

SC viability was assessed using the LIVE/DEAD assay (Ethidium homodimer-1, EthD-1; Thermo Fisher) at indicated timepoints. EthD-1-positive (dead) cells were quantified by fluorescence microscopy and expressed as a percentage of total DAPI-stained nuclei. For the MOI/post-infection timing assessment (Figure 3), 12-13 random images were analyzed per condition per biological replicate; for the 5-day aTc induction viability assessment (Figure 4), 4-5 random images were analyzed per condition per biological replicate (n = 3 independent experiments for each).

### 2.6 Nano-Luciferase (NLuc) Reporter Assay

At indicated timepoints following aTc addition, SCs were lysed using Nano-Glo lysis buffer (Promega). NLuc activity was measured on a VarioSkan Lux Microplate reader (Thermo Scientific) using the Nano-Glo Luciferase Assay System per manufacturer instructions. Luminescence (in relative luminescence units, RLU) was normalized either to intracellular colony forming units (CFUs) determined prior to aTc induction, or to SC number, as indicated in each figure legend.

### 2.7 Bacterial Clearance — Colony Forming Unit Assay

To quantify residual intracellular *Lm*, SCs were washed with PBS, lysed in 0.1% Triton X-100 in sterile water, and serial dilutions of the lysate were plated on BHI agar plates. CFUs were counted after 48 h of incubation at 37 °C and expressed as CFU per well.

### 2.8 AlamarBlue assay

SC metabolic activity was assessed using the alamarBlue assay (cat. no. DAL1100, Thermo Fisher) as an indirect indicator of cell viability, based on the reduction of blue resazurin to pink resorufin by metabolically active cells. SCs were seeded at 60,000 cells per well in 96-well plates to establish a confluent monolayer.

Three conditions were included: untreated SCs maintained in normal growth medium; uninfected SCs treated with aTc and moxifloxacin; and SCs infected with *Lm* and subsequently treated with aTc and moxifloxacin. aTc was added at 7 h post-infection (hpi) and moxifloxacin at 13 hpi. Uninfected controls received both compounds at the corresponding experimental times.

Four plates were prepared per independent experiment, each assessed at a separate endpoint, as indicated in Table 2. Two plates were assessed after 36 or 48 h of moxifloxacin treatment, corresponding to 49 and 61 hpi. For the remaining two plates, treatment medium was replaced with normal growth medium at 49 or 61 hpi, followed by a 24 h recovery period, and these were assessed at 73 and 85 hpi, respectively.

**Table. 2.**
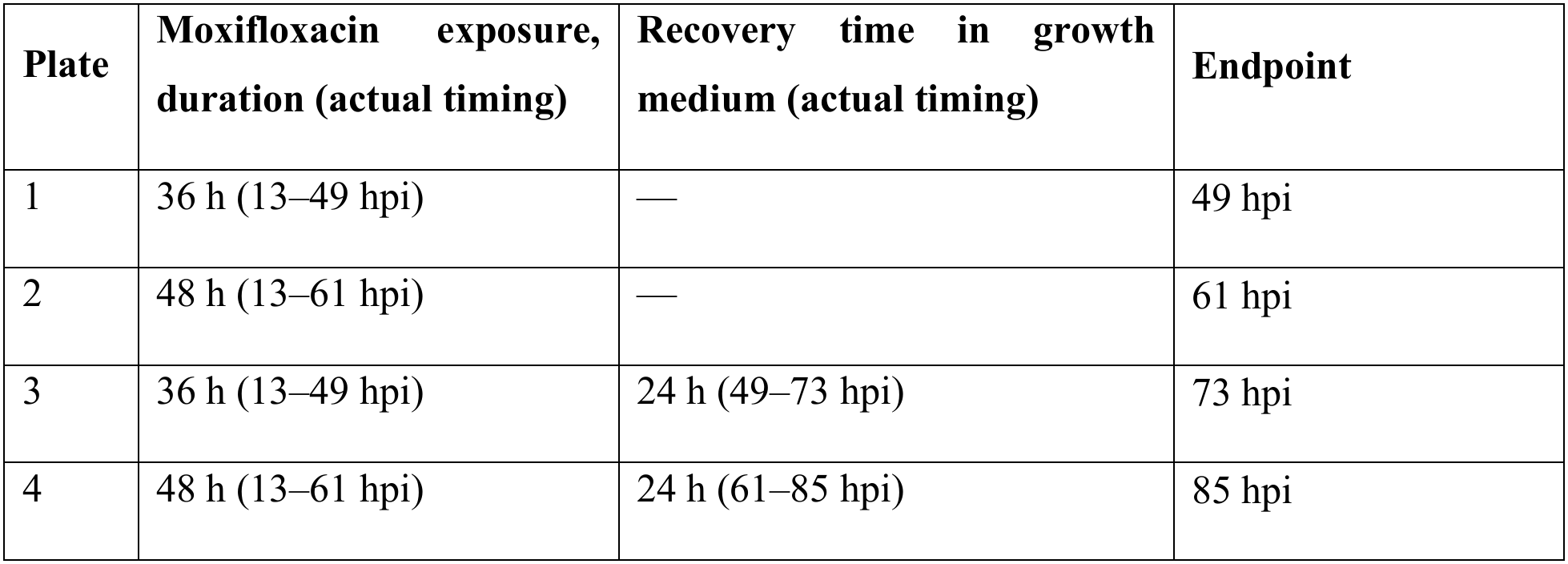
Transfection efficiencies of nonviral delivery methods in primary SCs.

At each endpoint, the culture medium in all wells was replaced with fresh growth medium containing 10% (v/v) alamarBlue. Plates were incubated for 4 h at 37 °C under 5% CO₂, after which absorbance was measured at 570 and 600 nm on a microplate reader. Percentage reduction of alamarBlue was calculated as:

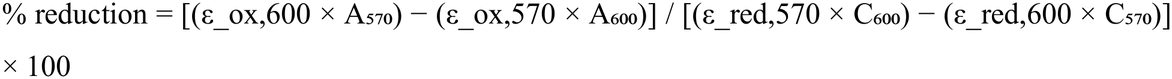

where A₅₇₀ and A₆₀₀ are sample-well absorbances, C₅₇₀ and C₆₀₀ are absorbances of cell-free control wells containing medium and alamarBlue, and ε_ox and ε_red are the molar extinction coefficients of the oxidized and reduced dye at the indicated wavelengths. Dual-wavelength measurement corrects for the overlapping absorbance spectra of the two dye forms.

Within each independent experiment, percentage reduction was normalized to the mean percentage reduction of untreated SC at the same endpoint:

Relative metabolic activity (%) = (% reduction_sample / mean % reduction_untreated, same endpoint) × 100 Untreated controls were thus assigned a relative metabolic activity of 100%. Three independent biological replicates were performed, each comprising five technical replicates per condition and endpoint; technical replicates were averaged to yield a single value per biological replicate.

### 2.9 Statistical Analysis

Statistical analyses were performed using GraphPad Prism 10. Data are presented as mean ± standard deviation (SD). Comparisons between two groups were performed using an unpaired two-tailed *t*-test. For experiments involving multiple groups, data were analyzed using one-way or two-way analysis of variance (ANOVA), followed by either Tukey’s or Dunnett’s multiple-comparisons test, as appropriate. A *P* value <0.05 was considered statistically significant.

## 3. Results

### 3.1 *Lm* Drones Enable Functional circRNA Bactofection of Primary Rat SCs

To establish whether *Lm* drones can support intracellular circRNA programming of primary SCs, we used the previously validated aTc-inducible TT-s split-NLuc circRNA delivery system [28] (Figure 1). Primary rat SC cultures were confirmed to be >95% pure by immunostaining for the SC-specific transcription factor SOX10 (Supplementary Figure 1). To characterize bacterial infection and intracellular localization, GFP-expressing *Lm* was added to SC cultures at an MOI of 20, and cells were examined 24 hpi. GFP-positive bacterial puncta were observed within SOX10-positive cells, consistent with intracellular bacterial localization (Figure 2A). Under these conditions, 94.4% of SCs were infected (Figure 2B).

**Figure 2.**
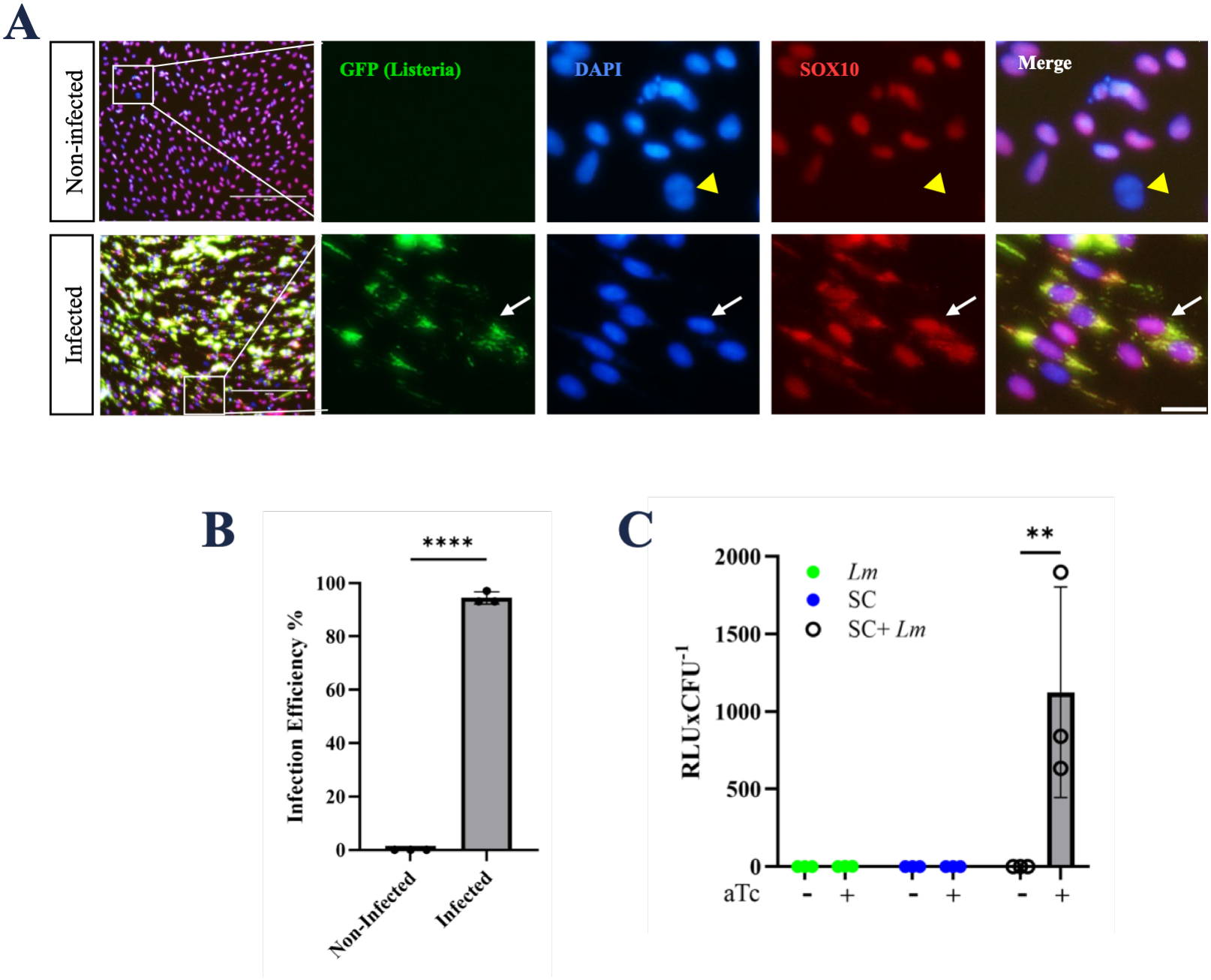
*Lm* Drones Enable Functional circRNA Bactofection of Primary SCs. **A.** Representative immunofluorescence images of noninfected (top) and *Lm*-infected (bottom) primary SC cultures showing GFP-expressing *Lm* (green), DAPI (blue), and the SC marker SOX10 (red). Enlarged inset images show GFP-positive intracellular *Lm* within SOX10-positive SCs (arrow). The arrowhead indicates a SOX10-negative cell, likely representing a contaminating fibroblast. Scale bars, 200 µm (main left images) and 20 µm (inset images). **B.** Quantification of infection efficiency in primary SCs. Data are presented as mean ± SD (n = 3). Statistical significance was determined using an unpaired two-tailed *t*-test. ****P < 0.0001. **C.** NLuc reporter activity normalized to intracellular bacterial burden determined prior to aTc induction. (RLU × CFU⁻¹) in *Lm* alone, non-infected SCs, and *Lm*-infected SCs (SC + *Lm*) with or without aTc induction. Significant NLuc activity above background was detected only in the SC + *Lm* + aTc condition. Data are presented as mean ± SD (n = 3). Statistical significance was determined using two-way ANOVA followed by Tukey’s multiple-comparisons test. **P < 0.01.

We next determined whether intracellular *Lm* could support functional circRNA delivery. NLuc activity was detected above background only in infected SCs treated with aTc (SC + *Lm* + aTc), whereas uninfected SCs and infected SCs without aTc induction showed only background activity (Figure 2C). The strict dependence of NLuc activity on both *Lm* infection and aTc induction establishes functional circRNA bactofection and protein expression in primary SCs.

### 3.2 Optimization of Infection Dose and Timing Defines an Operating Window for circRNA Bactofection

To establish operating conditions suitable for intracellular engineering of primary SCs, we systematically varied multiplicity of infection (MOI 5, 10, 20, 30, 40, 50, and 100) and quantified SC death (EthD-1 staining) and NLuc reporter activity at 4, 6, 8, and 24 h post-infection (Figure 3A, B; Supplementary Figure 2). During the first 8 hpi, SC viability remained high across the entire MOI range, with cell death remaining below 2% even at the highest MOI tested, MOI 100 (Figure 3A). In parallel, NLuc activity increased with increasing MOI during these early time points (Figure 3B). Extending the post-infection period to 24 h substantially altered the balance between reporter expression and cell viability. SC death increased across the MOI range, rising from 4% at MOI 5 to 29% at MOI 100 (Figure 3A), while NLuc activity also increased significantly (Figure 3B). Thus, increased reporter expression at 24 h was accompanied by markedly greater cytotoxicity.

**Figure 3.**
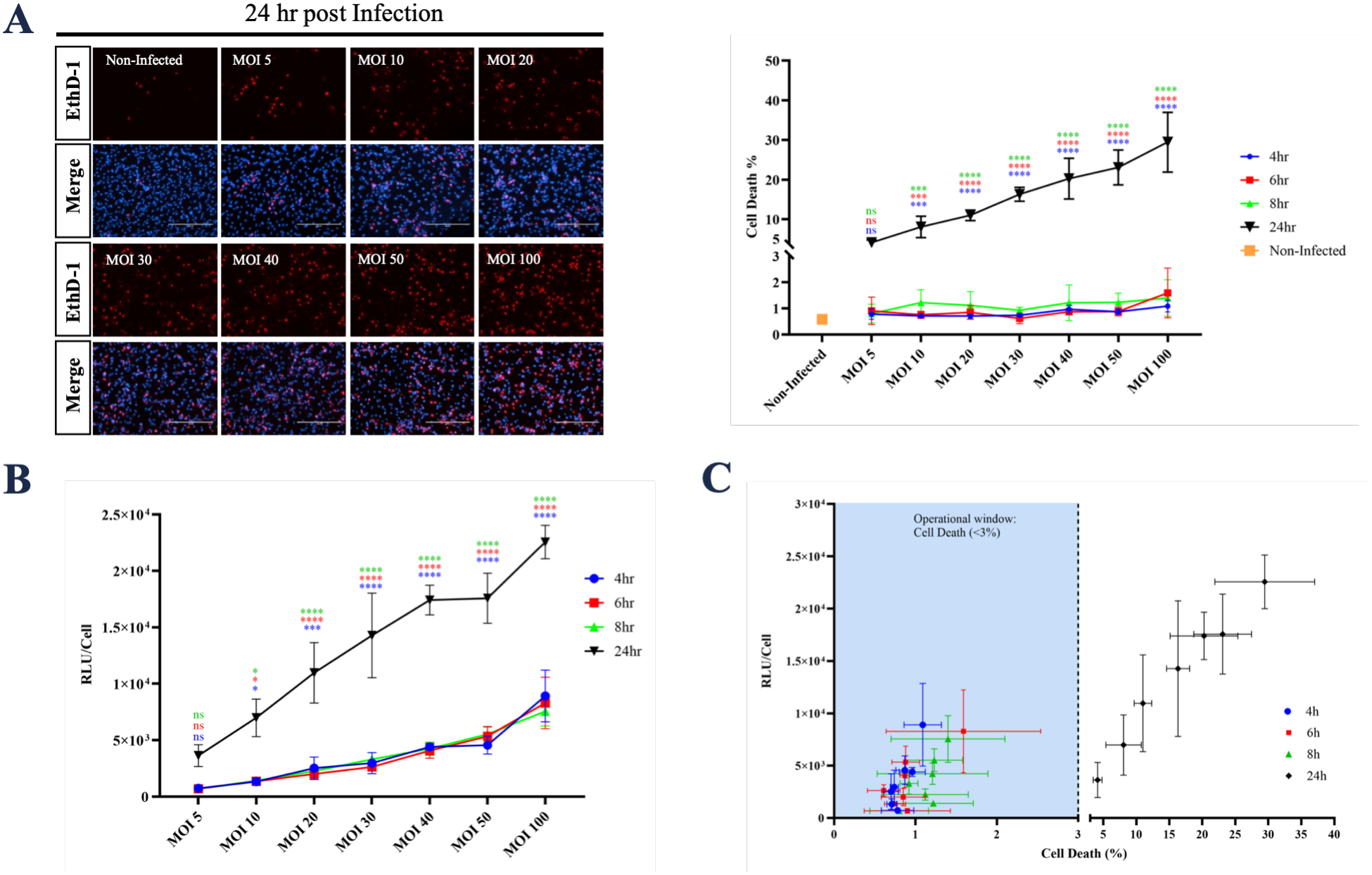
Optimization of Infection Dose and Timing Defines an Operational Window for circRNA Bactofection. **A.** Representative EthD-1/DAPI images of SCs infected with *Lm* across MOIs 5–100 at 24 hpi and quantification of cell death at 4, 6, 8, and 24 hpi. Cell death remained low at 4–8 h but increased substantially by 24 h, particularly at higher MOIs. Representative images for the 4, 6, and 8 h time points are shown in Supplementary Figure 2. Scale bar, 200 µm. **B.** NLuc reporter activity measured across the same MOI and post-infection time points. Reporter activity increased with increasing MOI and was highest at 24 h, which was accompanied by increased cytotoxicity. **C.** Scatter plot of NLuc reporter activity versus SC death across all MOI and post-infection time-point combinations tested. The blue shaded region indicates the defined operational window of <3% cell death, illustrating the range of conditions that maintain low cytotoxicity while supporting reporter expression. Data are presented as mean ± SD (n = 3). Statistical significance was determined by two-way ANOVA followed by Dunnett’s multiple-comparisons test. \**P* < 0.05, **P < 0.01, \*\*\**P* < 0.001, ****P < 0.0001; ns, not significant.

To define a practical operating window, NLuc activity was plotted against cell death across all MOIs and post-infection time points (Figure 3C). Because no conditions produced intermediate levels of cytotoxicity (2–4% cell death), we predefined <3% cell death as the threshold for acceptable toxicity. Under this criterion, the 4–8 h time points across the tested MOI range maintained low cytotoxicity while supporting detectable NLuc activity, whereas most 24 h conditions exceeded the threshold. Subsequent experiments were therefore performed within the early post-infection window, using MOI 10 and MOI 20 as representative conditions. Collectively, these experiments identified operating conditions that maximize circRNA delivery while preserving primary SC viability.

### 3.3 Transient circRNA Expression Supports Protein Production Without Sustained Cytotoxicity

Having established the operating conditions for circRNA bactofection, we next characterized the kinetics of inducible circRNA expression and its effect on SC viability over five days. A five-day observation window was selected to assess sustained circRNA expression while minimizing potential confounding effects associated with prolonged culture and subsequent cell splitting or replating. SCs bactofected at MOI 10 or MOI 20 were treated with aTc beginning 7 hpi and assessed daily for NLuc activity and EthD-1 staining (Figure 4). NLuc activity peaked one day after induction and progressively declined over the subsequent five days at both MOIs (Figure 4A). By day 5, mean NLuc activity had decreased approximately 3.2-fold relative to day 1 under both conditions, consistent with transient circRNA-mediated protein expression.

**Figure 4.**
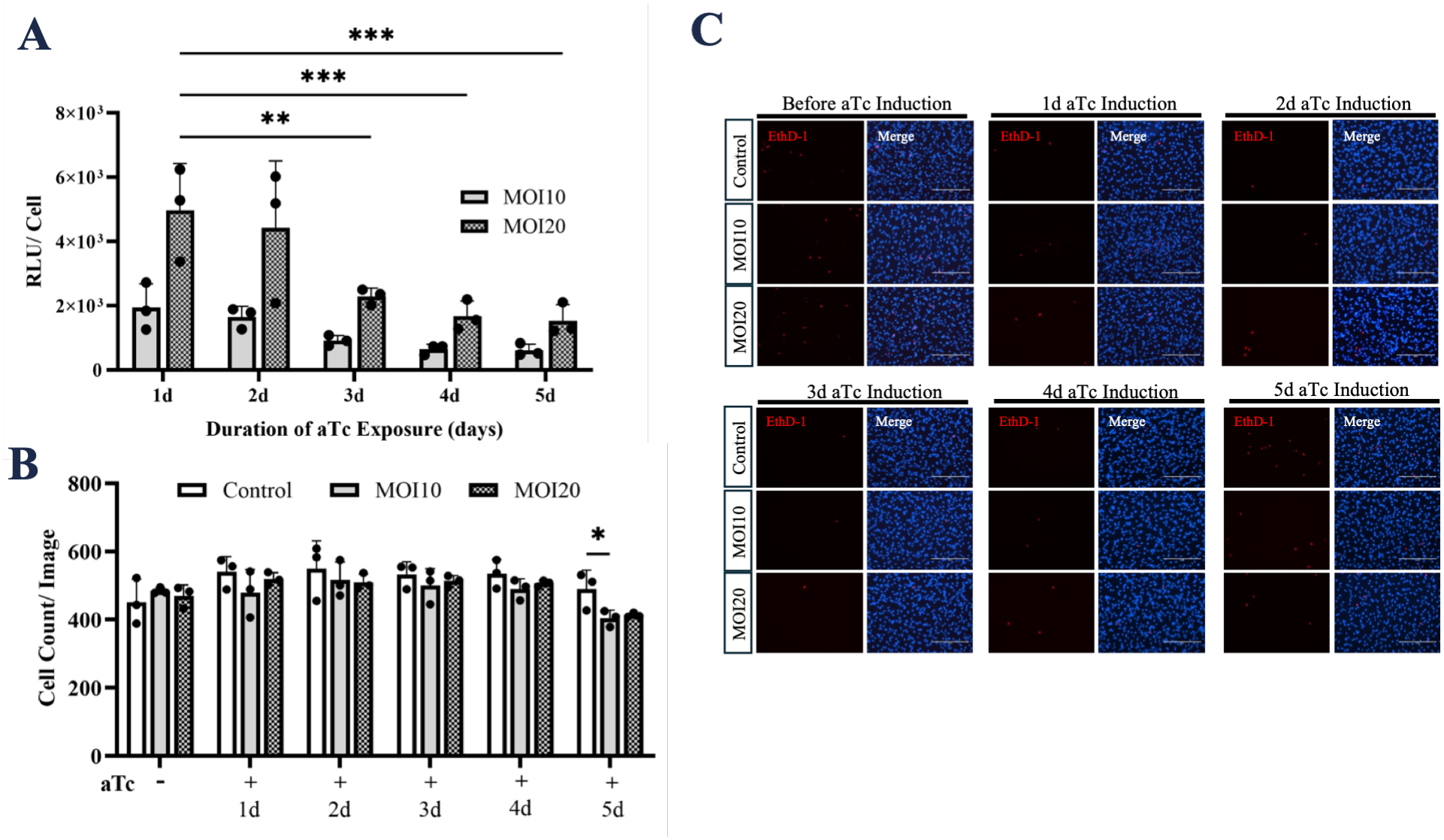
Transient circRNA Expression Supports Protein Production Without Sustained Cytotoxicity. **A.** NLuc reporter activity (RLU per cell) in SCs bactofected at MOI 10 or MOI 20, measured daily over 5 days of aTc induction. Reporter activity peaked on day 1 and progressively declined thereafter at both MOIs. **B.** Cell counts per imaging field in noninfected control, MOI 10-, and MOI 20-bactofected SCs over the 5-day aTc induction period. Cell numbers remained comparable across groups through day 4, with a modest reduction in infected cultures relative to control at day 5. **C.** Representative EthD-1/DAPI images (top) and quantification of cell death (bottom) in control, MOI 10-, and MOI 20-bactofected SCs before aTc induction and over the subsequent 5 days. SC death remained low throughout the induction period, with the only significant difference observed between control and MOI 10 before aTc induction. Data are presented as mean ± SD (n = 3). Statistical significance was determined by two-way ANOVA followed by Dunnett’s multiple-comparisons test. \**P* < 0.05, **P < 0.01, \*\*\**P* < 0.001. Scale bar, 200 µm.

Cell numbers per imaging field remained comparable among control, MOI 10, and MOI 20 cultures during the first four days of induction, with only a modest but statistically significant reduction observed on day 5 in bactofected cultures (Figure 4B). Consistent with these findings, EthD-1 staining remained low (≤1% for most conditions and time points) throughout the observation period (Figure 4C), indicating minimal cytotoxicity under the tested bactofection conditions. Together, these results demonstrate that transient circRNA expression can be maintained for several days without loss of SC viability.

### 3.4 Moxifloxacin Efficiently Eliminates Intracellular *Lm* Following circRNA Delivery

To determine whether intracellular bacteria could be eliminated after circRNA delivery, SCs bactofected at an MOI of 20 were treated with aTc from 7 to 13 h hpi. Cultures were subsequently maintained in moxifloxacin. Intracellular bacterial persistence was quantified 6 h after the start of aTc induction and at 24, 36, and 48 h after the initiation of moxifloxacin treatment (Figure 5A), corresponding to 13, 37, 49, and 61 hpi.

**Figure 5.**
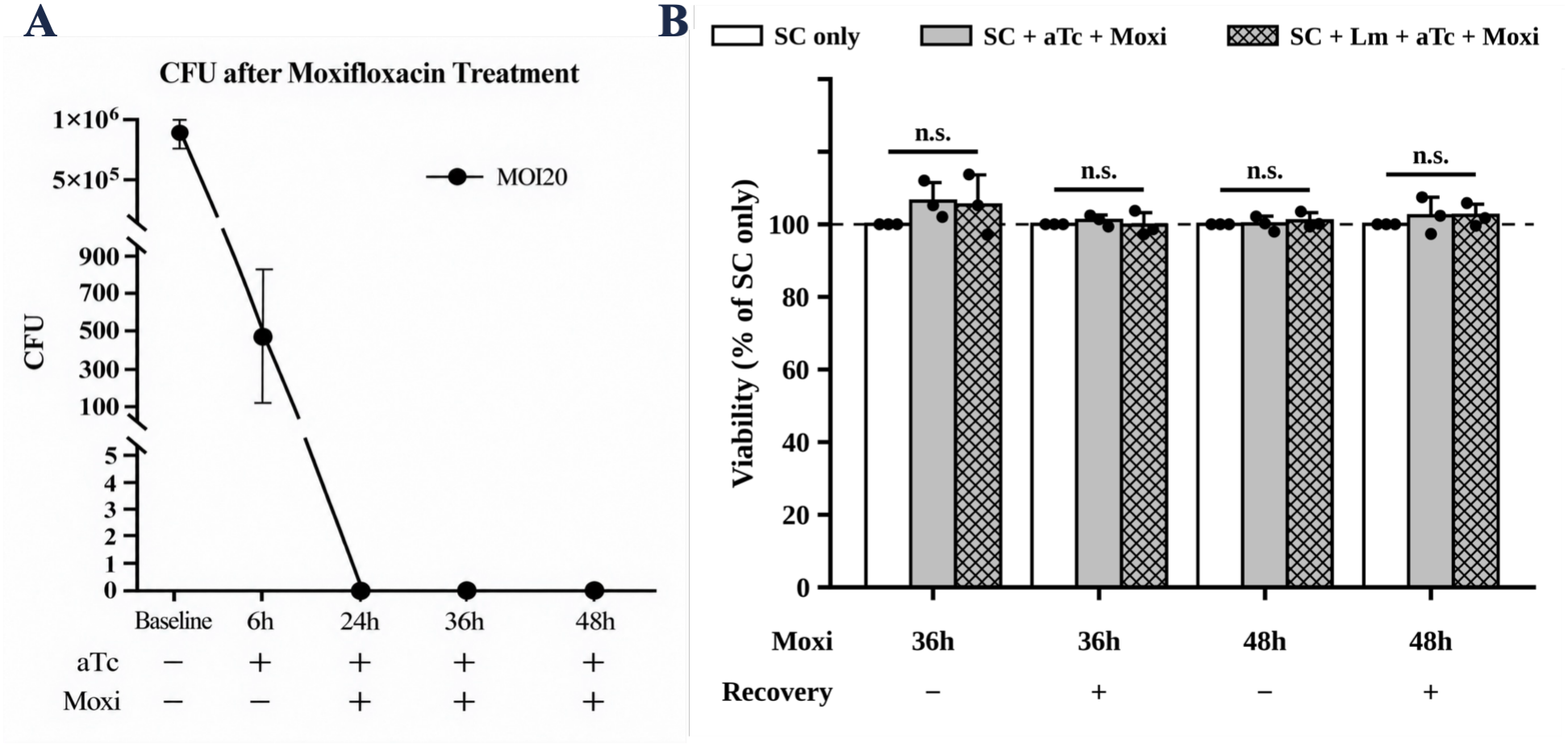
Moxifloxacin treatment eliminates intracellular *Lm* without significantly altering SC metabolic activity. **A.** Intracellular *Lm* burden in infected SCs at MOI 20, quantified as colony-forming units (CFU). Following the baseline measurement, cultures underwent a 6-h induction period with aTc (1 µg/mL), followed by moxifloxacin (4 µg/mL) treatment. Intracellular CFU decreased during the induction period and fell below the assay detection limit after 24 h of moxifloxacin treatment, remaining undetectable at 36 and 48 h. Data are presented as mean ± SD (n = 3). Statistical significance was determined by one-way ANOVA followed by Dunnett’s multiple-comparisons test. \*\**P* < 0.01; ns, not significant; \*\*\*\**P* < 0.0001. **B.** Relative SC metabolic activity measured using the alamarBlue assay after 36 or 48 h of moxifloxacin treatment, either immediately after treatment or following a 24 h recovery period in normal growth medium. Experimental groups comprised untreated SC (white bars), uninfected SC treated with aTc and moxifloxacin (grey bars), and *Lm*-infected SC treated with aTc and moxifloxacin (hatched bars). At each endpoint, culture medium was replaced with fresh medium containing 10% (v/v) alamarBlue, and cells were incubated for 4 h at 37 °C under 5% CO₂ before absorbance was measured at 570 and 600 nm. Percentage dye reduction was calculated using the molar extinction coefficients of the oxidized and reduced reagent and normalized to untreated SC at the corresponding endpoint, defined as 100% (dashed line). This readout reflects relative metabolic activity as an indirect indicator of cell viability. No statistically significant differences were detected for the comparisons indicated. Data are presented as mean ± SD (n = 3). Two-sided paired t-tests, run separately at each of the four timepoints without multiple-comparison correction; individual points represent biological replicate means. n.s., not significant.

In the absence of moxifloxacin, residual unlysed intracellular *Lm* persisted beyond the induction period. With moxifloxacin, bacterial burden was reduced to undetectable levels 24 h post-treatment and remained undetectable thereafter (Figure 5A). Moxifloxacin treatment therefore reliably clears residual intracellular *Lm* within 24 h of circRNA delivery under the tested conditions.

### 3.5 Bactofection Does Not Significantly Change SC metabolic activity

To assess whether bactofection compromises SC health, we compared the metabolic activity of untreated SC in normal growth medium, uninfected SC treated with aTc and moxifloxacin, and SC infected with *Lm* and subsequently treated with aTc and moxifloxacin (Figure 5B). SC were maintained in medium supplemented with forskolin and NRG1 and therefore remained in a proliferative, non-myelinating state throughout the experiment. Metabolic activity was measured by alamarBlue assay and normalized to untreated SC at each endpoint.

Cells were assessed after 36 or 48 h of moxifloxacin treatment, with or without an additional 24 h recovery in normal growth medium. At every endpoint, relative metabolic activity remained at approximately 100% of untreated controls in both treated groups (36 h: 106 ± 5% and 105 ± 8%; 36 h + recovery: 101 ± 1.5% and 100 ± 3%; 48 h: 100 ± 2% and 101 ± 2%; 48 h + recovery: 102 ± 5% and 102 ± 3% for SC + aTc + Moxi and SC + *Lm* + aTc + Moxi, respectively; mean ± SD, n = 3). No significant differences were detected between any group and the untreated control at any time point.

Overall, SC metabolic activity remained comparable to untreated controls following antibiotic treatment alone or bactofection followed by antibiotic treatment, with no significant differences detected at any assessed time point.

## Discussion

### 4.1 Principal Findings and Significance

This study establishes the first example of programmable intracellular engineering of primary cells using *Lm*-synthesized circRNA. Extending an attenuated, aTc-inducible *Lm* drone platform previously validated in immortalized cell lines, we show that engineered *Lm* drones invade primary rat SCs, synthesize and release functional circRNA, and support inducible protein expression while maintaining high SC viability under optimized conditions. Four operating parameters define the platform in this primary glial cell type: (i) efficient infection (>94% of SCs), (ii) strict aTc-dependent expression of a functional circRNA reporter, (iii) a broad early post-infection window in which SC death remained below 2% across a range of MOIs while supporting robust reporter expression, and (iv) complete elimination of intracellular *Lm* within 24 h after moxifloxacin treatment. These results establish an optimized workflow for transient circRNA programming of primary SCs (Figure 6) and demonstrate that engineered *Lm* drones can serve as programmable intracellular factories for therapeutically relevant primary cells.

**Figure 6.**
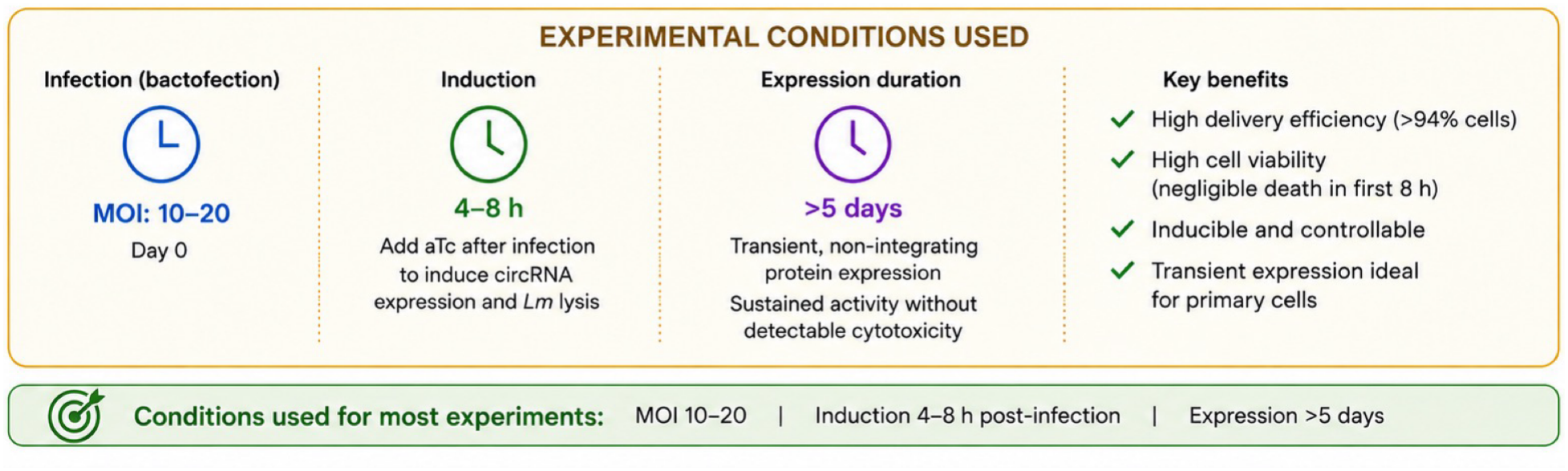
Summary of the *Lm*-mediated circRNA delivery strategy in primary SCs.

### 4.2 A Distinct Strategy for Primary Cell Engineering

Efficient engineering of primary cells remains a central challenge in various cell-based therapies. Viral vectors, lipid nanoparticles, electroporation, and nucleofection each have strengths, yet they differ in efficiency, cargo capacity, expression persistence, and effects on viability. The *Lm* drone platform offers a distinct paradigm that achieves higher efficiency than other methods for programming primary SCs (Table 1). Engineered bacteria invade target cells, synthesize circRNA under inducible control, circularize transcripts before release, and undergo programmed lysis to deliver translationally competent circRNA directly into the cytosol. Because only the circRNA cassette must be exchanged, the same bacterial chassis can be adapted to diverse protein payloads.

Although demonstrated here in SCs, the approach is intended as a general transient engineering platform whose adaptation to other difficult-to-transfect primary cell types should require only optimization of infection conditions.

The modular architecture also supports future multiplex programming. Delivery of multiple circRNA cassettes within a single bacterial chassis could enable coordinated expression of multiple proteins, an approach that remains challenging for many non-viral systems because each additional RNA species increases formulation complexity. Multiplex circRNA delivery was not tested here, but the platform’s design should facilitate combinatorial genetic programs in primary cells.

### 4.3 Operating Parameters for Transient circRNA Programming

Systematic variation of MOI and post-infection timing identified a practical operating window. Reporter activity increased with MOI but was strongly influenced by the duration of bacterial residence. During the first 4–8 hpi, robust expression was achieved while cell death remained below 2%. Extending the infection period to 24 h produced marked cytotoxicity. Thus, the duration of intracellular bacterial persistence is a critical determinant of the balance between engineering efficiency and cell viability.

The transient reporter profile further illustrates the behavior of the delivered circRNA. NLuc activity peaked one day after induction and gradually declined over the following five days despite minimal loss of viable SCs. Because bacterial lysis provides a discrete cargo-release event, this pattern is consistent with transient circRNA-mediated expression rather than continuous intracellular production. Direct measurements of circRNA abundance will be needed to distinguish RNA turnover from changes in translational activity. The broad low-toxicity window identified here should facilitate adaptation to additional primary cell types that require different infection conditions.

### 4.4 Limitations and Future Directions

Several limitations remain. All experiments were performed in monolayer cultures of primary rat SCs; performance in neuron–SC co-cultures, three-dimensional models, or injured peripheral nerve has not been examined. Only a single circRNA reporter was tested, so the effects of therapeutic payload size, protein complexity, or simultaneous delivery of multiple circRNAs remain unknown. Population-averaged measurements also do not resolve cell-to-cell variability in programming or expression. Longer-term effects on SC phenotype, proliferation, myelination capacity, and regenerative function remain to be determined.

Innate immune responses to the bacterial chassis or the circRNA cargo were not examined. Although circRNAs exhibit significantly reduced immunogenicity relative to linear RNAs, lysed *Lm* can activate multiple host sensing pathways. Defining how bacterial invasion, circRNA production, and programmed clearance influence inflammatory signaling will be essential for therapeutic applications.

Despite these limitations, the work establishes a methodological foundation for programmable intracellular engineering of primary cells using bacterially synthesized and intracellularly delivered circRNA. The modular platform should enable rapid exchange of payloads and adaptation to other therapeutically relevant primary cell types, including immune, epithelial, mesenchymal, and progenitor cells. These studies therefore position self-destructing *Lm* drones not merely as an alternative delivery vehicle, but as a versatile intracellular engineering platform for transient, non-integrating protein expression in primary cells, particularly those, such as SCs, whose regenerative functions can be enhanced by time-limited expression of therapeutic factors.

## 5. Conclusions

This study establishes the first demonstration of programmable intracellular engineering of primary SCs using bacterially synthesized circRNA. Extending a previously validated self-destructing, aTc-inducible *Lm* drone platform from immortalized cell lines to primary cells, we show efficient bacterial invasion, inducible functional expression of circRNA cargo, and an optimized early post-infection window that supports transient protein expression while maintaining high SC viability. Under the conditions tested (MOI 10–20), circRNA-mediated protein expression was achieved in >94% of primary SCs, persisted for several days with minimal cytotoxicity, and intracellular bacteria were completely eliminated within 24 h by moxifloxacin treatment.

These findings define infection dose and timing as key determinants of the balance between engineering efficiency and cell viability, and they establish an optimized workflow for *Lm*-mediated circRNA programming of primary SCs. Although further work is needed to evaluate immune responses, long-term effects on SC phenotype and regenerative function, and performance with therapeutic or multiplex payloads, the platform provides a foundation for transient, non-integrating protein expression in difficult-to-transfect primary cells and complements existing RNA and viral delivery technologies.

## SUPPLEMENTARY MATERIALS

**Supplementary Figure 1.**
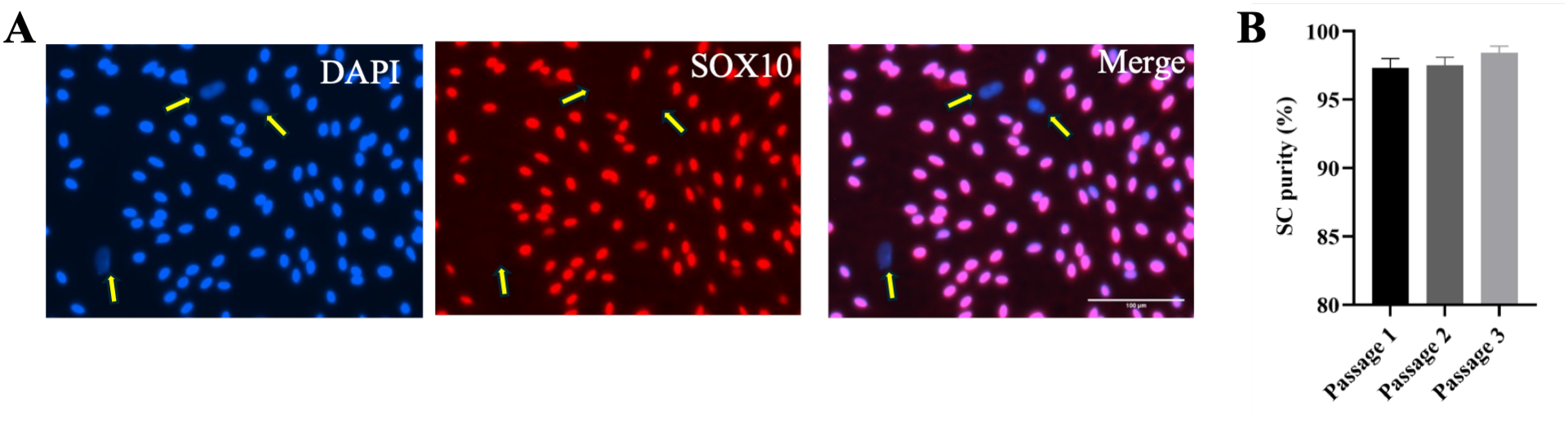
Purity assessment of primary rat SC cultures. **A.** Representative immunofluorescence images of primary rat SCs stained for SOX10 (red) and DAPI (blue). The merged image shows predominantly SOX10+/DAPI+ SCs, with rare SOX10-negative nuclei (yellow arrows), likely representing fibroblasts. Scale bar, 200 µm. **B.** Quantification of SC purity, defined as the percentage of SOX10+ cells among total DAPI+ nuclei, across three sequential passages (P1–P3). SC purity remained >95% across all passages. Data are presented as mean ± SD (n = 3).

**Supplementary Figure 2.**
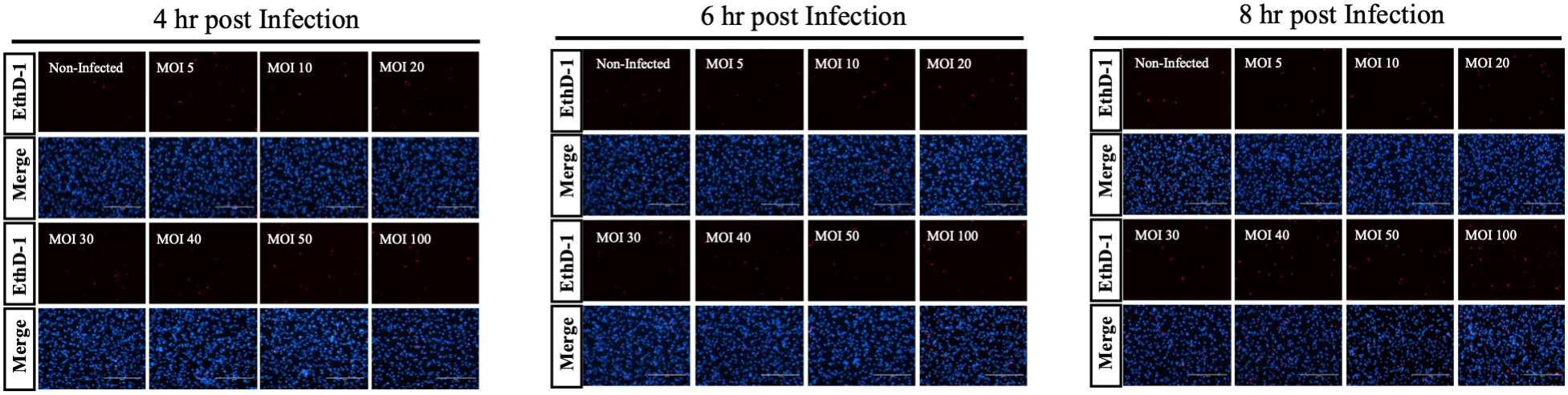
Representative EthD-1 staining of SCs across MOI and early post-infection timepoints. EthD-1 (dead cell, red)/DAPI merge images of SCs at 4, 6, and 8 hr after infection with *Lm* across MOIs of 5–100. Minimal EthD-1 positivity is observed across all MOIs at these early timepoints, consistent with the quantification in Figure 2a. n = 3 independent experiments per condition. Scale bar, 200 µm.

